# Profiling G protein-coupled receptors of *Fasciola hepatica* identifies orphan rhodopsins unique to phylum Platyhelminthes

**DOI:** 10.1101/207316

**Authors:** Paul McVeigh, Erin McCammick, Paul McCusker, Duncan Wells, Jane Hodgkinson, Steve Paterson, Angela Mousley, Nikki J. Marks, Aaron G. Maule

## Abstract

G protein-coupled receptors (GPCRs) are established drug targets. Despite their considerable appeal as targets for next-generation anthelmintics, poor understanding of their diversity and function in parasitic helminths has thwarted progress towards GPCR-targeted anti-parasite drugs. This study facilitates GPCR research in the liver fluke, *Fasciola hepatica*, by generating the first profile of GPCRs from the *F. hepatica* genome. Our dataset describes 146 high confidence GPCRs, representing the largest cohort of GPCRs, and the most complete set of *in silico* ligand-receptor predictions, yet reported in any parasitic helminth. All GPCRs fall within the established GRAFS nomenclature; comprising three glutamate, 135 rhodopsin, two adhesion, five frizzled and one smoothened GPCR. Stringent annotation pipelines identified 18 highly diverged rhodopsins in *F. hepatica* that maintained core rhodopsin signatures, but lacked significant similarity with non-flatworm sequences, providing a new sub-group of potential flukicide targets. These facilitated identification of a larger cohort of 76 related sequences from available flatworm genomes, representing new members of existing groups of flatworm-specific rhodopsins. These receptors imply flatworm specific GPCR functions, and/or co-evolution with unique flatworm ligands, and could facilitate development of exquisitely selective anthelminthics. Ligand binding domain sequence conservation relative to deorphanised rhodopsins enabled high confidence ligand-receptor matching of seventeen receptors activated by acetylcholine, neuropeptide F/Y, octopamine or serotonin. RNA-Seq analyses showed expression of 101 GPCRs across various developmental stages, with the majority expressed most highly in the pathogenic intra-mammalian juvenile parasites. These data identify a broad complement of GPCRs in *F. hepatica*, including rhodopsins likely to have key functions in neuromuscular control and sensory perception, as well as frizzled and adhesion families implicated, in other species, in growth, development and reproduction. This catalogue of liver fluke GPCRs provides a platform for new avenues into our understanding of flatworm biology and anthelmintic discovery.

**Author Summary:** *Fasciola* spp. liver fluke are important veterinary pathogens with impacts on human and animal health, and food security, around the world. Liver fluke have developed resistance to most of the drugs used to treat them (flukicides). Since no vaccines exist, we need to develop new flukicides as a matter of urgency. Most anthelmintic drugs used to treat parasitic worm infections operate by impeding the functioning of their nerve and muscle. In flatworms, most nervous signals are received by a type of receptor called a G protein-coupled receptor (GPCR). Since GPCRs control important parasite functions (e.g. movement, egg-laying, feeding), they represent appealing targets for new flukicides, but have not yet been targeted as such. This work exploited the *F. hepatica* genome to determine the quantity and diversity of GPCRs in liver fluke. We found more GPCRs in the *Fasciola* genome than have been reported in any other parasitic worm. These findings provide a foundation that for researchers to determine the functions of these receptors, and which molecules/ligands they are activated by. These data will pave the way to exploring the potential of *F. hepatica* GPCRs as targets for new flukicides.

## Introduction

*Fasciola spp*. liver fluke are pathogens of veterinary ruminants that threaten the sustainability of global meat and dairy production. Infection with *Fasciola* (fasciolosis/fascioliasis) inhibits animal productivity through liver condemnation, reduced meat and milk yields, and reduced fertility (for recent impact surveys see [1–4]. *Fasciola* spp. also infect humans, with fascioliasis considered a neglected tropical disease [5]. Anthelmintic chemotherapy currently carries the burden of fluke control, since there are no liver fluke vaccines [6]. Six flukicidal active compounds are available for general use, with on-farm resistance reported for all except oxyclozanide [7]. Resistance to the frontline flukicide, triclabendazole, also exists in human *F. hepatica* infections [8,9]. Given the absence of alternative control methods, new flukicides are essential for secure future treatment of veterinary and medical liver fluke infections.

The helminth neuromuscular system is a prime source of molecular targets for new anthelmintics [10–12], not least because many existing anthelmintics (dichlorvos, levamisole, morantel, piperazine, pyrantel, macrocyclic lactones, paraherquamide, amino acetonitrile derivatives) act upon receptors or enzymes associated with classical neurotransmission in nematodes [11] The G protein-coupled receptors (GPCRs) that transduce signals from both peptidergic and classical neurotransmitters are of broad importance to helminth neuromuscular function. Despite industry efforts to exploit helminth GPCRs in the context of anthelmintic discovery [13], only a single current anthelmintic (emodepside) has been attributed GPCR-directed activity as part of its mode of action [14–16]. GPCRs are druggable targets, since 33% of human prescription medicines are attributed a GPCR-based mode of action [17].

Despite two *F. hepatica* genomes [18,19], no GPCR sequences have been reported from *F. hepatica*. In contrast, GPCRs have been profiled in the genomes of trematodes (*Schistosoma mansoni* and *Schistosoma haematobium* [20,21]), cestodes (*Echinococcus multilocularis, E. granulosus, Taenia solium* and *Hymenolepis microstoma* [22]), and planaria (*Schmidtea mediterranea, Girardia tigrina* [21–24]). These datasets illustrated clear differences in the GPCR complements of individual flatworm classes and species, with reduced complements in parasitic flatworms compared to planarians.

This study profiles the GPCR complement of the temperate liver fluke *F. hepatica* for the first time, permitting comparisons with previously characterised species that inform evolutionarily and functionally conserved elements of flatworm GPCR signalling. We have identified and classified 146 GPCRs by GRAFS family (glutamate, rhodopsin, adhesion, frizzled, secretin) assignment [25], the majority of which are expressed in *Fasciola* RNA-Seq datasets. These include clear orthologues of GPCRs activated by known neurotransmitters, within which we performed the deepest *in silico* ligand-receptor matching analyses to date for any parasitic helminth. The latter predicted ligands for 17 *F. hepatica* GPCRs, designating these as primary targets for deorphanisation. Intriguingly, the dataset included a set of flatworm-expanded GPCRs lacking orthologues outside of phylum Platyhelminthes. Evolution of such GPCRs across the parasitic flatworm classes may have been driven by flatworm-specific functional requirements or co-evolution with flatworm ligands, either of which could help support novel anthelmintic discovery. This dataset provides the first description of GPCRs in liver fluke, laying a foundation for future advances in GPCR-directed functional genomics and flukicide discovery.

## Results and Discussion

### A first look at GPCRs in the *F. hepatica* genome

This study represents the first description of the GPCR complement of the temperate liver fluke, *F. hepatica*. Using HMM-led methods to examine available *F. hepatica* genome datasets, we identified 166 GPCR-like sequences in *F. hepatica* (Figure 1, S1 Table). Figure 1B shows that 49.7% contained 7 TM domains, with 88% of sequences containing at least four TMs. The remainder of this manuscript focuses on 146 sequences containing ≥4TM domains (S1 Table; S2 Text). Twenty sequences containing ≤3 TMs were analysed no further (Figure 1).

**Fig 1:**
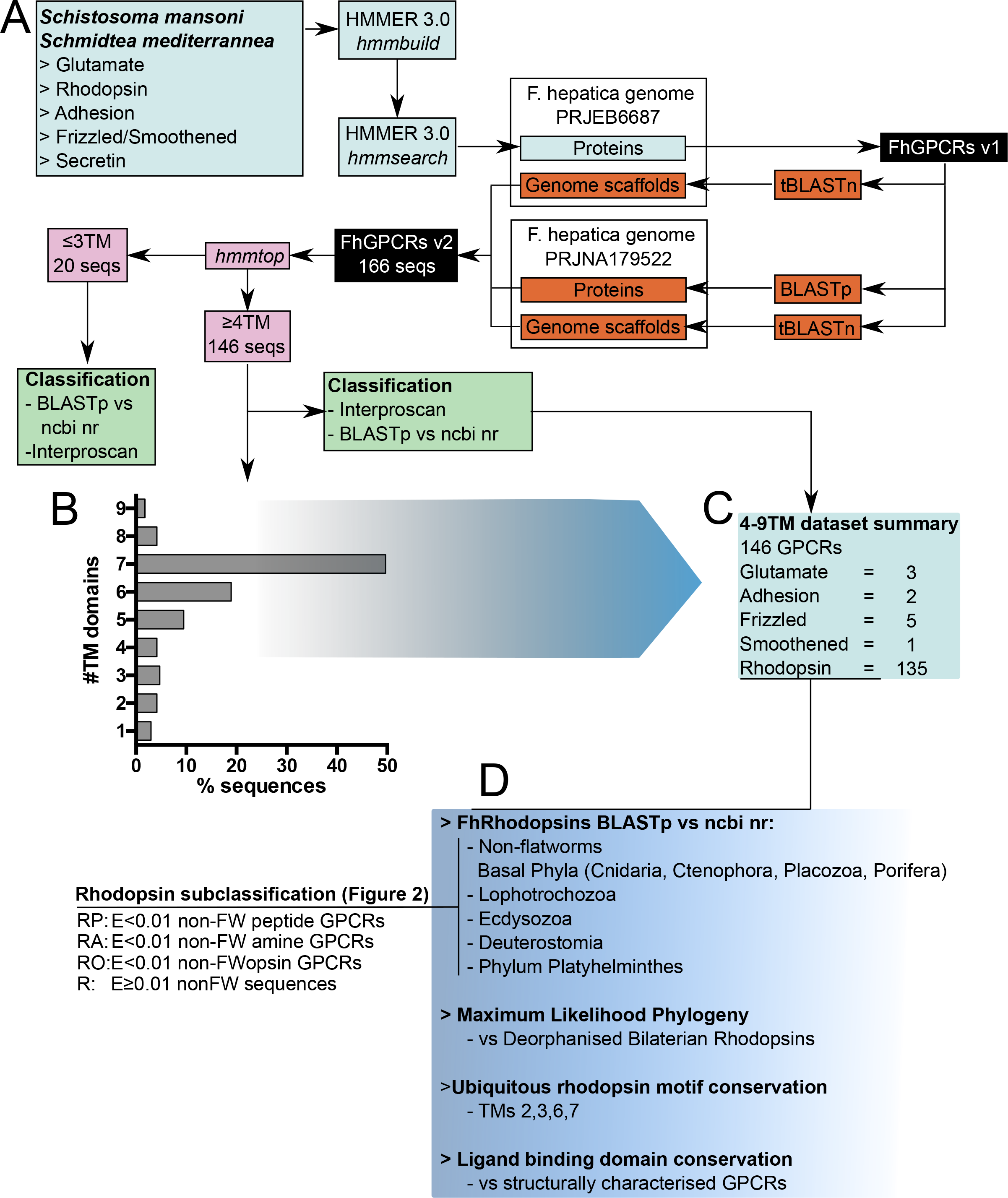
Methods for discovery and annotation of *Fasciola hepatica* G protein coupled receptors (FhGPCRs). (A) Hidden Markov Models (HMMs) representing glutamate, rhodopsin, adhesion, frizzled/smoothened and secretin families, and two rhodopsin subfamilies, were built from protein multiple sequence alignments of *Schistosoma mansoni* and *Schmidtea mediterranea* GPCRs [21]. HMMs were built and searched respectively using the *hmmbuild* and *hmmsearch* modules of HMMER v3.0. Searches were performed against two publically available *F. hepatica* genomes using *hmmsearch* and basic local alignment search tool (BLAST) tools. Each putative FhGPCR sequence was assessed for transmembrane (TM) domain composition with *hmmtop* before classification using tools including BLASTp, Interproscan and CLANS. (B) The largest proportion (49%) of FhGPCRs carried the full complement of 7 TMs, with 88% of sequences bearing at least 4 TMs. (C) GRAFS composition of 146 FhGPCRs carrying ≥4 TMs. (D) Rhodopsins were subject to further classification, including BLASTp vs datasets representing major non-flatworm animal phyla and superphyla. These rhodopsin homology classifications fed back into phylogenetic analyses versus deorphanised bilaterian GPCRs to confirm their putative ligand selectivity, with a final analysis of ligand binding domain composition comparing conservation of ligand interacting residues for characterised GPCRs reported in the literature with our *F. hepatica* assignments.

Our ≥4TM dataset (146 sequences) was comprised of three glutamate, 135 rhodopsin, two adhesion, five frizzled, and one smoothened GPCR. Sequence coverage was generally good in terms of TM and extracellular domain representation, so we did not attempt to extend truncated sequences into full-length receptors. The overall dataset contained excellent representation of seven TM domains, while N-terminal extracellular LBDs and cysteine-rich domains (CRD) were also detected (in glutamate, frizzled/smoothened, adhesion families). However, we could not identify N-terminal secretory signal peptides in any sequence, suggesting incomplete sequence coverage at extreme N-termini. Rhodopsins are designated by ubiquitously conserved motifs on TMs 2, 3, 6 and 7. All rhodopsin sequences contained at least one of these motifs (Figure 2, S3 Table), including in the highly diverged flatworm-specific rhodopsins described below.

**Fig 2:**
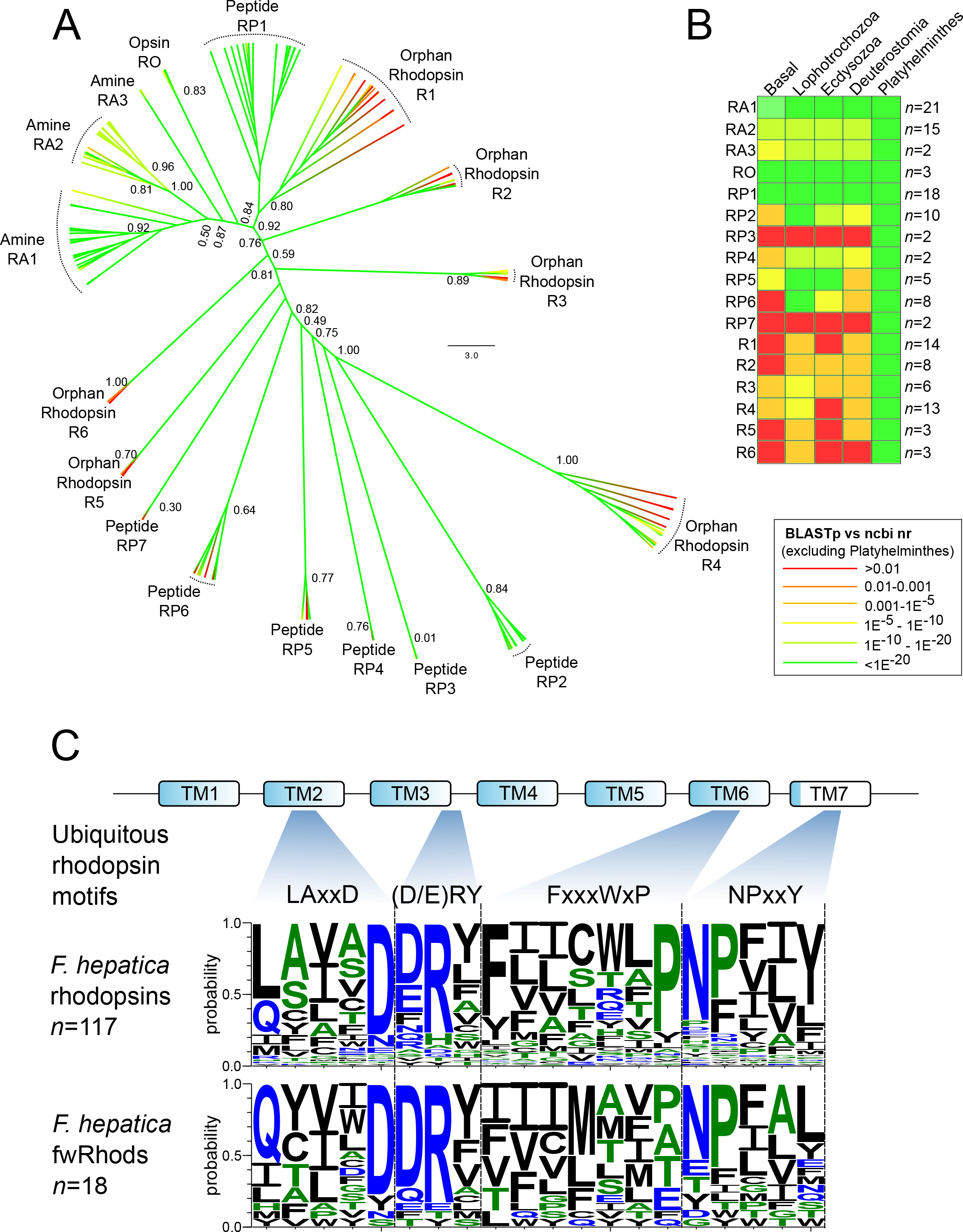
Phylogenetic classification of *Fasciola hepatica* rhodopsin G protein-coupled receptors. (A) Maximum-likelihood cladogram of *F. hepatica* rhodopsins. Phylogeny delineated clades containing rhodopsins with distinct homologies (RA, amine; RP, peptide; RO, opsin: R, orphan rhodopsin). The orphan clades contained sequences with generally low BLASTp similarity to their closest non-flatworm BLASTp hit, but concentrated within them were 18 sequences with exceptionally low (E>0.01) BLASTp similarity to non-flatworm sequences (fwRhods). The tree was midpoint rooted and was generated from a multiple protein sequence alignment trimmed to TM domains I-VII. Numbers at nodes indicate statistical support from approximate likelihood ratio test (aLRT). Tip colours are coded according to the E-value scale (as indicated) of that GPCR’s closest BLASTp match in the ncbi nr database, excluding phylum Platyhelminthes. (B) Summary of sequence similarity comparisons between GPCRs within each rhodopsin clade, and their closest BLASTp hits in major phylogenetic groups (Basal: Cnidaria, Ctenophora, Porifera, Placozoa; superphylum Lophotrochozoa, omitting Platyhelminthes; Superphylum Ecdysozoa; superphylum Deuterostomia; phylum Platyhelminthes). BLASTp E-value (median) is summarised in each case, colour coded as a heat map on the same colour scale as (A). The number of GPCRs comprising each *F. hepatica* clade (*n*) is also indicated. (C) Sequence diversity within ubiquitous rhodopsin motifs of the majority (117) of the *F. hepatica* rhodopsins (upper panel), compared to those motifs in 18 *F. hepatica* fwRhods (lower panel). The mammalian consensus motifs are illustrated above the top panel, along with an illustration of the location of each motif within the rhodopsin 7TM domain structure. Some variability is visible within the TM2 and TM6 motifs, but TM3 and TM7 motifs are well conserved.

Table 1 compares the *F. hepatica* GPCR complement with other flatworms, illustrating that *F. hepatica* has the largest GPCR complement reported from any parasitic flatworm to date. The bulk of the expansion involves rhodopsins, while the other GRAFS families are comparable between *F. hepatica* and other flatworm parasites. Secretin is the notable exception, at least one of which has been identified in every other species studied, but which was absent from the datasets scrutinized here.

**Table 1.** Comparison of the *Fasciola hepatica* G-protein coupled receptor (GPCR) complement with those reported from other flatworms. Species complements are shown in the context of GRAFS nomenclature [25]. * Saberi et al [24] described 566 GPCRs in *Schmidtea mediterrannea*, of which 516 fall within GRAFS nomenclature.

|  | *F. hepatica* | *S. mansoni*<sup>[21]</sup> | *S. mansoni*<sup>[20]</sup> | *S. haematobium*<sup>[20]</sup> | *E. multilocularis*<sup>[22]</sup> | *S. mediterranea*<sup>[21]</sup> | *S. mediterranea*<sup>[24]</sup> |
| --- | --- | --- | --- | --- | --- | --- | --- |
| Glutamate | 3 | 2 | 2 | 2 | 5 | 9 | 11 |
| Rhodopsin | 135 | 105 | 59 | 53 | 48 | 418 | 461 |
| Adhesion | 2 | 3 | - | - | 4 | 9 | 14 |
| Frizzled/Smoothed | 6 | 5 | 4 | 4 | 5 | 11 | 10 |
| Secretin | 0 | 2 | 5 | 5 | 1 | 1 | 20 |
| **Total** | **146** | **117** | **64** | **64** | **83** | **448** | **516\*** |

### Stringent annotation of flatworm-specific orphan rhodopsin GPCRs in *F. hepatica*

Encompassing 135 sequences, the rhodopsin family is the largest of the GRAFS classifications in *F. hepatica*. Rhodopsins comprise four subfamilies (α, β, γ and δ) [26]; we identified members of both α and β groups, with nucleotide-activated (P2Y) receptors (γ group), and olfactory (δ group) receptors absent from our dataset (Figures 1, 2; S1 Table). The *F. hepatica* α subfamily contained 38 amine receptors and three opsins, with the β subfamily comprised of at least 47 peptide receptors. Homology-based annotations were supported by an ML phylogeny (Figure 2A), which clearly delineated between amine and opsin α clades, and the peptide-activated β-rhodopsin clades. Amine and peptide receptors were further delineated by additional phylogenetic and structural analyses, permitting high-confidence assignment of putative ligands to 16 GPCRs (see below).

Six clades contain an additional 44 rhodopsin sequences with low scoring (median E = 5.6e^−5^) similarity matches to a range of disparate α and β rhodopsins. Due to the subsequent difficulty in designating these clades as amine, peptide or opsin, we labelled them orphan rhodopsins (“R” clades in Figure 2A). Eighteen GPCRs within the orphan clades displayed exceptionally low similarity scores relative to non-flatworm sequences (Figure 2A,B). Seven returned no-significant hits in BLASTp searches against non-flatworm members of the ncbi nr dataset (the most diverse sequence dataset available to the research community), and the remaining eleven scored E>0.01. Domain analysis (InterPro) identified rhodopsin domains (IPR000276 or IPR019430) in thirteen of these (S1 Table, S3 Table), confirming their identity as rhodopsin-like GPCRs. More troublesome to classify were five that, in addition to lacking significant BLASTp identity to non-flatworm sequences, also lacked any identifiable protein domains/motifs (with the exception of TM domains). We annotated these as rhodopsins because: (i) They did not contain motifs/domains representative of any other protein family; (ii) They displayed topological similarity to GPCRs (ten had seven TM domains, seven had six TMs, one had five TM domains); (iii) They contained at least two of the conserved rhodopsin motifs in TM domains 2, 3, 6 and 7 similar to those seen in the rest of the *F. hepatica* rhodopsins (Figure 2C; S4 Table). As highly diverged rhodopsins with little or no sequence similarity versus host species, these 18 *F. hepatica* receptors have obvious appeal as potential targets for flukicidal compounds with exquisite selectivity for parasite receptors over those of the host. This potential is contingent on future work demonstrating essential functionality for these receptors; showing their wider expression across flatworm parasites would enable consideration of anthelmintics with multi-species activity. To investigate the latter question, we used BLASTp to search the 18 *F. hepatica* rhodopsins against other available genomes representing phylum Platyhelminthes.

### An orphan family of lineage-expanded rhodopsins in flatworm genomes

Although lacking similarity against non-flatworm datasets, each of the 18 lineage-expanded *F. hepatica* rhodopsins returned high-scoring hits in BLASTp searches against the genomes of other flatworms (WormBase Parasite release WBPS9). All returns were subsequently filtered through a stringent five-step pipeline (Figure 3A) consisting of: (i) Removal of duplicate sequences; (ii) Exclusion of sequences containing fewer than four TM domains; (iii) A requirement for reciprocal BLASTp against the *F. hepatica* genome to return a top hit scoring E<0.001 to one of the original 18 *F. hepatica* queries; (iv) A requirement for BLASTp against ncbi nr non flatworm sequences to return a top hit scoring E>0.01; (v) Removal of sequences lacking conservation of the ubiquitous rhodopsin motifs seen in the divergent *F. hepatica* rhodopsins (Figures 2C, 3C). The latter motifs were largely absent from cestode rhodopsins (with the exception of a single sequence from *Diphyllobothrium latum*, and three sequences from *Schistocephalus solidus*), and present in only two sequences from a single monogenean (*Protopolystoma xenopodis*). This left our final dataset consisting of 76 “flatworm-specific” rhodopsins (fwRhods; Figure 3B, Table S4) in phylum Platyhelminthes, heavily biased towards trematodes (70 sequences). Nineteen sequences from nine species of cestode were omitted from the final dataset despite meeting the inclusion criteria in most respects, because they lacked conservation of ubiquitous rhodopsin motifs (filtering step (v)). Although their further characterisation was beyond the scope of this study, they warrant more detailed examination in future studies as potential cestode-specific rhodopsins. Note that our filtering pipeline also excluded initial hits from *Gyrodactylus salaris* (Monogenea), and the Turbellarians *Macrostomum lignano* and *S. mediterranea*. Individual species complements of fwRhods showed some consistency (Figure 3B); the trematodes *F. hepatica* and *Echinostoma caproni* (both phylum Platyhelminthes Order Echinostomida) bore 18 and 19 sequences, respectively, most species of family Schistosomatidae contained 3-4 sequences each. The inclusion of two cestode species and a single monogenean may be an indication of the existence of distantly related rhodopsins in those lineages, rather than a true measure of the extent of cestode and monogenean fwRhod diversity. Again, proper classification of these groups will require further more focused study that was beyond the scope of the current work.

**Fig 3:**
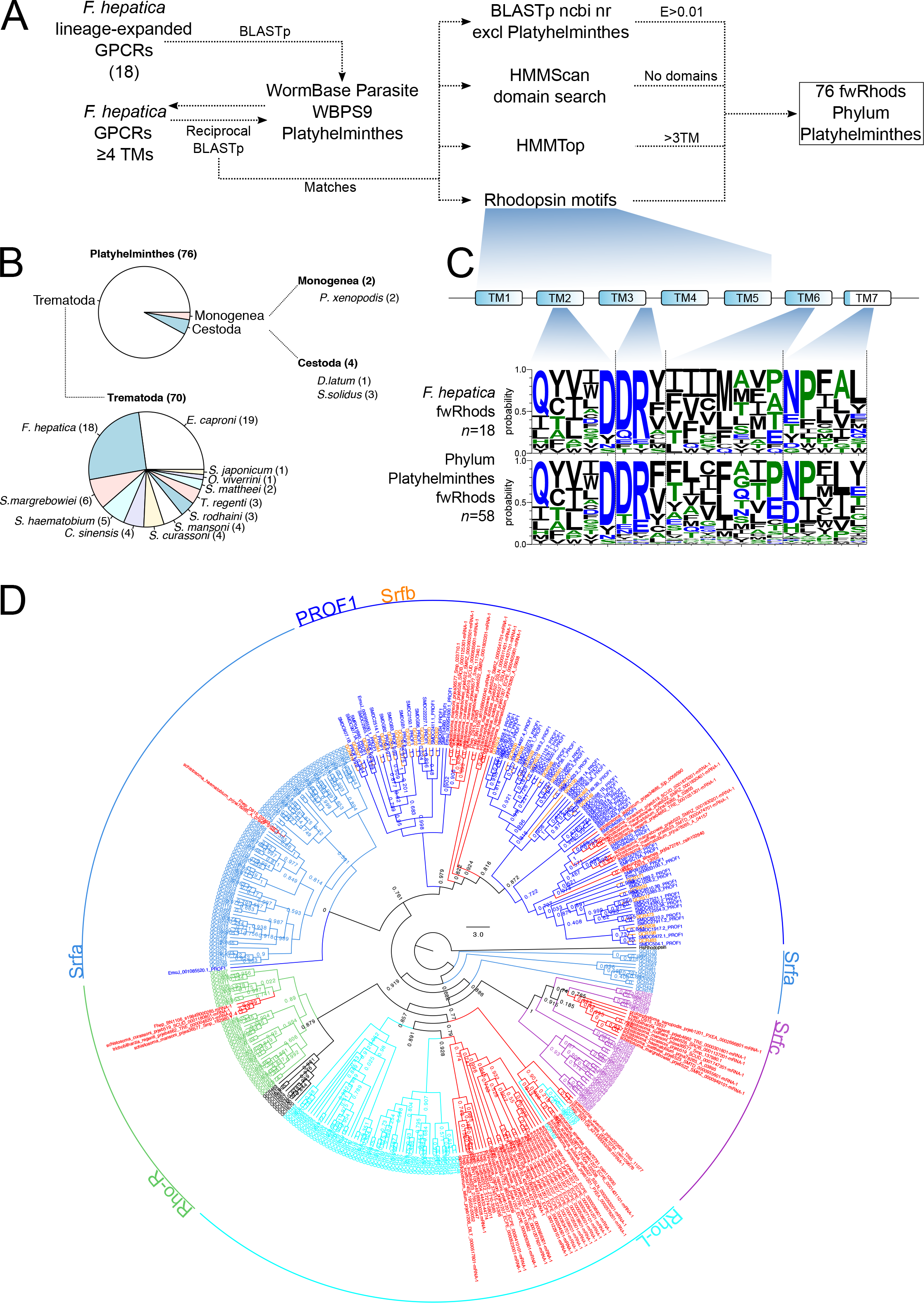
Identification of flatworm-specific rhodopsins (fwRhods) in genomes from phylum Platyhelminthes. (A) The 18 *Fasciola hepatica* GPCRs in our dataset that had poor BLASTp similarity (E>0.01) to non-flatworm sequences in the ncbi nr dataset (lsGPCRs), were used as queries in BLASTp searches of flatworm genomes in WormBase Parasite (release WBPS9). All hits scoring E<0.01 were back-searched by BLASTp against our *F. hepatica* GPCR dataset. Sequences scoring E<0.01 against one of the original *F. hepatica* GPCRs were retained as matches. These sequences were then filtered to identify those lacking matches in ncbi nr, lacking non-GPCR protein domains, possessing at least 4 transmembrane (TM) domains, and containing rhodopsin motifs consistent with those seen in the majority of *F. hepatica* rhodopsins (see C). (B) This process identified 76 fwRhods in phylum Platyhelminthes, the majority (70) of which were from class Trematoda. Small numbers were returned from classes Cestoda and Monogenea. Note that no fwRhods fitting these criteria were identified in class Turbellaria. (C) Sequence diversity within ubiquitous rhodopsin motifs of 18 *F. hepatica* fwRhods (upper panel), compared to those motifs in the 58 fwRhods identified in the wider phylum (lower panel); motifs are broadly similar between *F. hepatica* and the rest of the phylum. (D) Maximum likelihood phylogeny of 76 fwRhods, alongside flatworm-specific rhodopsins described previously (70 platyhelminth rhodopsin orphan family 1 (PROF1) [21,22], and 245 *S. mediterranea* G protein coupled receptor [GCRs, comprising RhoL, RhoR, Srfa, Srfb and Srfc families, reported as lacking non-flatworm homologues [24]) with branches coloured to indicate Family (dark blue, PROF1; mid blue, Srfa; cyan, Rho-L; green, Rho-R; orange, Srfb; purple, Srfc; red, fwRhod). Tree was rooted to a human rhodopsin (P08100) and was generated from an alignment trimmed to transmembrane domains I-VII. Numbers at nodes indicate statistical support from approximate likelihood ratio test (aLRT).

Our method for identification of fwRhods is supported by a similar BLAST-driven approach used to identify highly diverged “hidden orthologues” in flatworms [27]. It should be noted that the existence of sequences lacking sequence similarity to genes of other species is not a new finding. “Taxonomically-restricted genes” comprise 10-20% of every sequenced eukaryote genome, and may be essential for phylum-specific morphological and molecular diversity [28]. How do our fwRhods compare to previously reported groups of flatworm restricted GPCRs in *S. mansoni, S. mediterrannea and E. granulosus* [21,22,24]? Phylogenetic comparisons of these groups (Figure 3D) demonstrated that the previously described *Schmidtea* Srfb cluster [24] and the PROF1 clade (*E. multilocularis, Schmidtea, S. mansoni* [21]) are equivalent, and likely represent a single group. Our phylogeny added 23 fwRhods to this clade, including three from *F. hepatica* (BN1106_s6156B000040, D915_03083, D915_13002). Figure 3D designated the remaining fwRhods within additional pre-existing groups [24], placing 34 within Rho-L (including eight from *F. hepatica*), nine in Srfc (one from *F. hepatica*), four in Rho-R (one from *F hepatica*) and two in Srfa (one from *F. hepatica*). Four fwRhod sequences were omitted from this tree due to poor alignment.

Our approach to classifying flatworm-restricted rhodopsins was to err towards stringency, and this may have resulted in erroneous exclusion of some sequences from the dataset. There is no set definition for lineage specificity in the literature, but ours is the most stringent yet applied to flatworm GPCRs. Applying our BLASTp E≥0.01 cutoff (modified from Pearson [29]) to the previously described groups of flatworm-specific rhodopsins [21,22,24], excludes 57 of the 62 PROF1s described from *S. mansoni* and *S. mediterranea* and 287 of the 318 RhoL/R and Srfa/b/c flatworm-specific clusters in *S. mediterranea*. Further pursuit of the extent of lineage restricted GPCRs in the wider phylum was beyond the scope of this study, but we are currently trawling for taxonomically restricted flatworm GPCRs on a phylum wide scale.

We have established the existence of a group of rhodopsin GPCRs that appear restricted to, and expanded in, phylum Platyhelminthes. By definition these receptors are orphan (i.e. their native ligands are unknown), so key experiments must focus on identifying their ligands and functions. Such experiments can exploit the expanding molecular toolbox for flatworm parasites, which in *F. hepatica* includes RNA interference (RNAi) [30–33], interfaced with enhanced in vitro maintenance methods, and motility, growth/development and survival assays [31,34,35]. Our phylogeny (Figure 2A) suggests that fwRhods are more similar to peptide than amine receptors. If their heterologous expression can be achieved, one approach to characterisation would be to screen them with the growing canon of peptide ligands from flatworms [36–38], as well as from other genera, in a receptor activation assay. Subsequent localisation of their spatial expression patterns would provide additional data that would inform function.

### Predicting ligands for *F. hepatica* rhodopsin GPCRs

In addition to the flatworm-specific fwRhod sequences described above, for which the ligands and functions remain cryptic, we also identified many rhodopsins with clear similarity to previously annotated GPCRs. Figure 2A shows the phylogenetic delineation of these sequences into amine-, opsin-and peptide-like receptors, distinctions that are supported by BLASTp comparisons with general (ncbi nr) and lineage-specific (superphylum level) datasets, as well as by gross domain structure (InterProScan) (S1 Table). These data provided a foundation for deeper classification of putative ligand-receptor matches.

The structure and function of GPCR LBDs can be studied using molecular modelling to predict interactions with receptor-bound ligands. These predictions can then be validated by targeted mutagenesis of residues within the LBD, measuring impacts with downstream signalling assays. Such experiments have been performed in model vertebrates and invertebrates, enabling identification of evolutionarily conserved binding residues/motifs. These data inform assignment of putative ligands to newly discovered receptors. Since mutagenesis experiments have not yet been performed in flatworm GPCRs, we employed a comparative approach to identify 17 *F. hepatica* rhodopsins with LBD motifs diagnostic of receptors for NPF/Y, 5-HT, octopamine (Oct) or acetylcholine (ACh) (Figure 4; S4 Table), thus enabling *in silico* ligand-receptor matching of these GPCRs.

**Fig 4:**
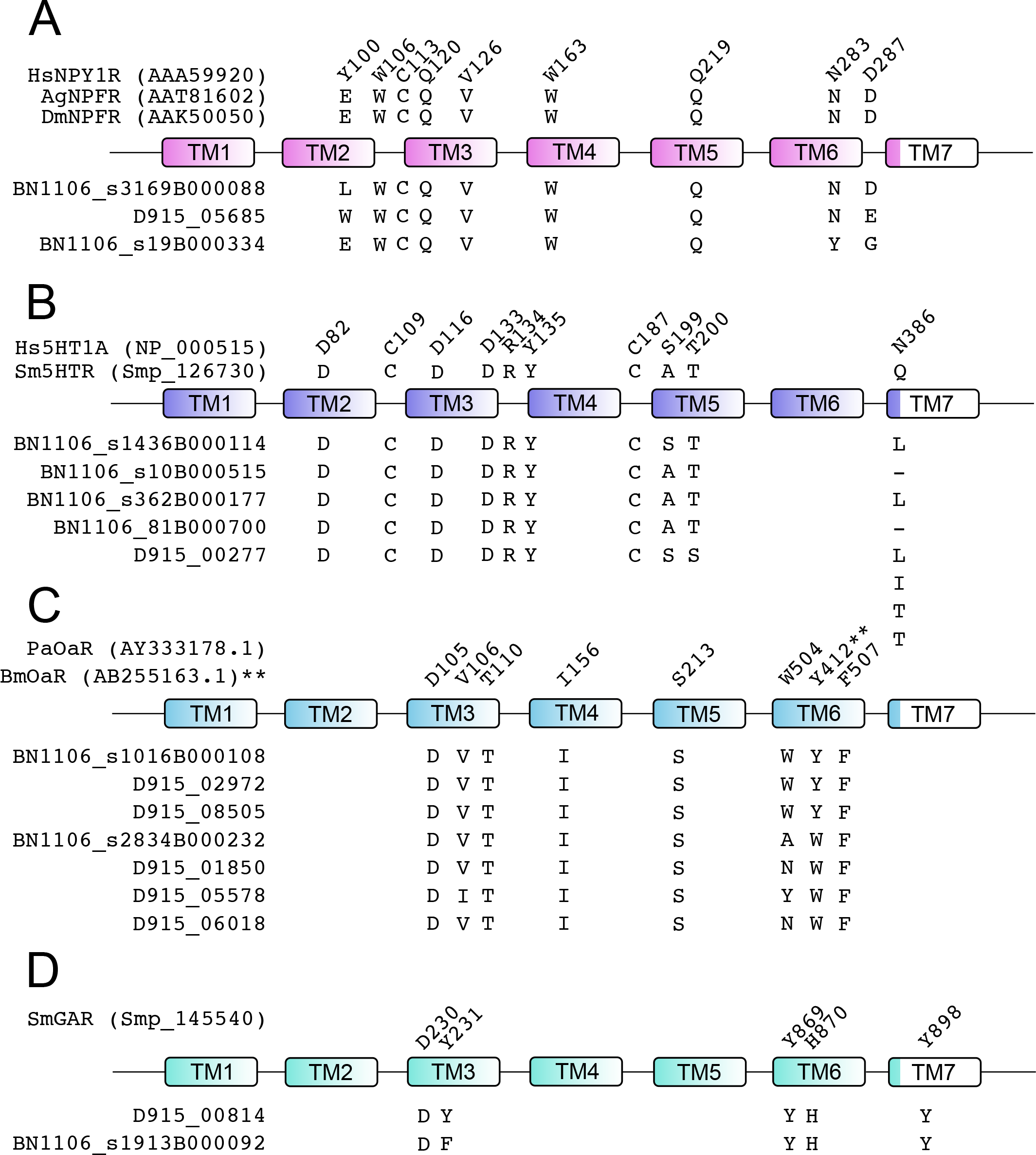
Conservation of ligand-interacting residues between 17 *Fasciola hepatica* G protein-coupled receptors (GPCRs) and structurally characterized homologues from other species. (A) Neuropeptide F/Y receptor ligand binding residues as characterised by mutagenesis in human neuropeptide Y receptor NPY1R [39–43], and conserved in *Anopheles gambiae* (Ag) and *Drosophila melanogaster* (Dm) neuropeptide F receptors (NPFR) [44]. Numbering relative to HsNPY1R. (B) Serotonin (5-hydroxytryptamine; 5HT) receptor ligand binding residues as characterised by mutagenesis in human 5HT receptor (Hs5HT1A) [102], and conserved in *Schistosoma mansoni* 5HTR [51]. Numbering relative to Hs5HT1A. (C) Octopamine receptor (OaR) ligand binding residues as characterised by homology modelling of the *Periplaneta americana* (Pa) [58], and mutational analysis of the *Bombyx mori* (Bm) [59] octopamine receptor ligand binding domain. Numbering relative to PaOAR, except for Y412 which is shown relative to BmOAR. (D) Acetylcholine receptor ligand binding residues as characterised by homology modelling of the *S. mansoni* G protein-coupled acetylcholine receptor (SmGAR) [62]; numbering relative to SmGAR. In each case, only *F. hepatica* sequences displaying at least 75% identity across the stated ligand binding residues are shown. Relative positions of residues across seven transmembrane domains (TM1-7) are shown. TM diagrams are not to scale.

Comparison of *F. hepatica* rhodopsins by structural alignment with LBD residues conserved across vertebrate NPY and dipteran NPF receptors [39–44] identified three peptide receptors with more than 75% identity across 9 ligand-interacting positions (Figure 4A). The two highest scoring GPCRs (BN1106_s3169B000088 and D915_05685) are also found, in our phylogenetic analysis (S5 Figure) in the same clade as the deorphanized NPF/Y receptors of human (HsNPYR2), *Glossina mortisans* (Glomo-NPFR) and *S. mediterranea* (SmedNPYR1). These data designate these three *F. hepatica* GPCRs as prime candidates for further work to deorphanize and confirm these receptors as NPF/Y-activated, and to probe the biology of NPF/Y receptors in parasitic flatworms. A single NPF/Y receptor has been functionally characterised in *S. mediterranea*, displaying a role in the maintenance of sexual maturity [24]. If related functions are conserved in liver fluke NPF/Y receptors they could have appeal as therapeutic targets in adult fluke that could interrupt parasite transmission, although their utility for the control of acute fasciolosis, caused by migrating juveniles, would be open to question.

Broad phylogenetic comparison of our peptide receptor set with a comprehensive collection of deorphanized bilaterian rhodopsin GPCRs (S5 Figure), identified *F. hepatica* receptors similar to those for myomodulin, FLP, luqin and Neuropeptide KY (NKY). These ligands have all been predicted or demonstrated in previous biochemical or in silico studies of flatworm neuropeptides [36–38]. We also uncovered *F. hepatica* GPCRs with phylogenetic similarity to allatotropin, allatostatin, thyrotropin-releasing hormone and sex peptide receptors. These ligands have not yet been reported in flatworms, although the existence of allatostatin-like receptors in flatworms is supported by the inter-phyla activity of arthoropod allatostatins in helminth (including flatworm) neuromuscular assays [45].

No *F. hepatica* neuropeptide sequences have been published yet, but our unpublished data suggest the presence of at least 36 neuropeptide genes in the *F. hepatica* genome (Duncan Wells, Queen’s University Belfast, personal communication). These ligands would facilitate deorphanisation of heterologously-expressed peptide GPCRs (S1 Table). This is essential work, since although two planarian peptide receptors have been deorphanised [23,24], no flatworm parasite peptide GPCRs have been ligand matched yet. Receptor deorphanisation provides a starting point for drug discovery, by enabling development of agonists or antagonists that modulate the interaction of a GPCR with its cognate ligand. Such compounds could form the basis of ligand series for screening pipelines that would lead to new flukicides [46,47].

Serotonin (5-hydroxytryptamine, 5-HT) is abundant throughout flatworm nervous systems, and is considered the primary flatworm excitatory neurotransmitter [48]. Deorphanized GPCRs activated by 5-HT have been described in turbellarians and trematodes, with an *S. mansoni* 5-HT receptor (Sm5HTR) involved in neuromuscular control [49–51]. Five *F. hepatica* rhodopsins (Figure 4B) bore appreciable (≥80%) positional identity in amino acids shown to be key ligand-interacting residues in the human 5HT1A LBD [52,53]. Notably, these residues were also conserved in the deorphanized *S. mansoni* 5-HT receptor (Sm5HTR, Smp_126730) [51]. Three of the sequences (BN1106_s362B000177, BN1106_s81B000700 and BN1106_s10B000515) also resembled Sm5HTR in our phylogenetic analysis, identifying them as likely 5-HT receptors. The remaining two (D915_00277 and BN1106_s1436B000114) appeared phylogenetically more similar to an *S. mansoni* dopamine receptor (Smp_127310) [54]. These annotations provide rational starting points for receptor deorphanization using functional genomic and/or heterologous expression tools. We found that *F. hepatica* dopamine-like receptors, identified by phylogeny (S5 Figure), displayed poor conservation (max 56% overall identity) to the human D2 LBD [55]. Due to this lack of selectivity, we did not annotate any *F. hepatica* GPCRs as dopamine receptors.

Although common in other invertebrates, octopamine has not yet been directly demonstrated as a neurotransmitter in flatworms. Evidence for its presence is indirect, based on tyramine β-hydroxylase (octopamine’s biosynthetic enzyme) activity in cestodes and planaria [56,57]. Three rhodopsins (Figure 4C) showed 100% conservation of the arthropod octopamine LBD, as determined from *Periplaneta americana* and *Bombyx mori* [58,59], with an additional four showing 88% conservation. Of these seven rhodopsins, four resolved in close phylogenetic proximity to *Drosophila* mushroom body octopamine receptors (D915_02972), *Drosophila* octopamine beta-receptors (D915_08505 and BN1106_s1016B000108) (S5 Figure) or a *Drosophila* tyramine receptor (D915_05578), denoting these as high-confidence octopamine receptors. These data provide further evidence in support of a functional role for this enigmatic classical neurotransmitter in flatworms.

Acetylcholine has species-specific impacts on flatworm neuromuscular preparations *in vitro*, with myoinhibitory effects in *Fasciola* [60]. Two putative muscarinic acetylcholine receptors (mAChRs), shared highest LBD identity with a Rat M3 ACh receptor (Figure 4D) [61]. Although these were only 67% identical to the rat sequence, the five ligand-interacting residues within their LBDs were 100% identical to those of a deorphanised *S. mansoni* mAChR, known to be involved in neuromuscular coordination (SmGAR) [62]. These receptors (D915_00814 and BN1106_s1913B000092) were also the most similar to SmGAR in our phylogeny (S5 Figure) so we consider them amongst our high confidence candidates for deorphanization.

### *F. hepatica* glutamate receptors bear divergent glutamate binding domains

At least three glutamate-like GPCRs exist in *F. hepatica* (Figure 5A, S1 Table). All three are defined by significant BLASTp similarity (median E=2.3e^−34^) to metabotropic glutamate receptors (mGluRs), and/or by the presence of InterPro GPCR family 3 (Class C) domains IPR017978, IPR000162 or IPR000337. Phylogenetic analysis of these GPCRs was performed alongside receptors representative of the various Class C subgroups (Figure 5) [63], including Ca^2+^-sensing receptors, γ-aminobutyric acid type B (GABA_B_) receptors, metabotropic glutamate (mGluR) receptors, and vertebrate taste receptors; for reference we also included the two previously reported mGluRs from *S. mansoni* [21]. One *F. hepatica* GPCR (BN1106_s2924B000081) resolved alongside Smp_052680, which has previously been described as an *S. mansoni* mGluR; these receptors form a close outgroup from the mGluR clade, supporting their designation as mGluRs. A second *F. hepatica* glutamate receptor (BN1106_s1717B000113) also has a close *S. mansoni* ortholog (Smp_128940), both of which reside in an orphan outgroup that is of uncertain provenance. The third *F. hepatica* glutamate receptor resides within another orphan group, close to human GPR158 and GPR179, two closely related class C GPCRs expressed respectively in the human brain and retina [64]. Although these receptors have been linked with specific disease states [65,66], their ligands remain unknown.

**Fig 5:**
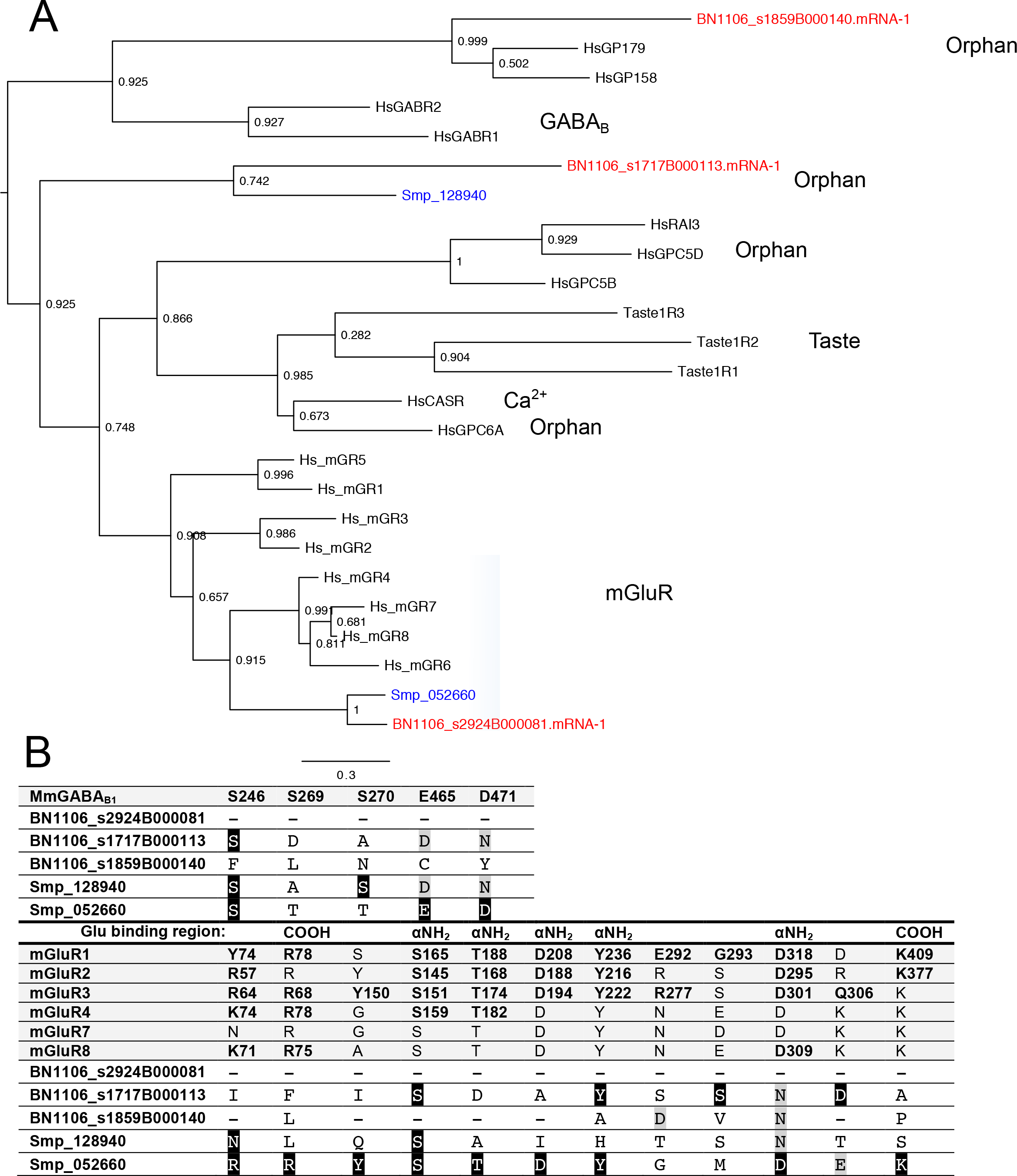
*Fasciola hepatica* glutamate G-protein coupled receptors (GPCRs) display divergent phylogeny and ligand binding domain (LBD) composition. (A) Maximum likelihood phylogeny containing three *F. hepatica* glutamate receptors, alongside representative receptors from the various recognised GPCR Class C subgroups (subclasses indicated by blue boxes: Ca^2+^, Ca^2+^-sensing receptor; GABA_B_, γ-aminobutyric acid type B receptors; mGluR, metabotropic glutamate receptors; Orphan, receptors with no known ligand; Taste, vertebrate taste receptors). Two previously reported *Schistosoma mansoni* glutamate receptors are also included; *F. hepatica* sequences are coloured red, *S. mansoni* are coloured blue, all others are black. Node numbers indicate statistical support as determined by approximate likelihood ratio test (aLRT). Tree was midpoint rooted. (B) Conservation of ligand-interacting residues between vertebrate GABA_B_ and metabotropic glutamate receptors (mGluR), and *F. hepatica* class C GPCRs. Agonist-interacting residues were identified by multiple protein sequence alignment of *F. hepatica* glutamate receptors against mutationally-identified ligand interacting residues (those causing a significant reduction in receptor signalling activity), from mouse GABA_B_ receptor (top panel), or selected human mGluR subtypes (lower panel). Identical amino acids in *F. hepatica/S. mansoni* GPCRs are represented by white text on black background, functionally conserved amino acids by black text on grey background. In lower panel, mutations causing a significant reduction in mGluR receptor activity are bold and numbered, with the region of the glutamate molecule bound by each residue indicated (COOH, C-terminus; NH_2_, N-terminus). For references see [63,103,104].

Divergence within the LBD can inform the ligand selectivity of Class C receptors [21,67]. To further classify the two orphan glutamate GPCRs described above, we generated multiple sequence alignments to analyse the conservation of established agonist-interacting residues between mammalian mGluR and GABA_B_ receptors and our *F. hepatica* GPCRs. These analyses identified no significant conservation of either mGluR or GABAB_B_ LBD residues (Figure 6B). Figure 6B also includes the previously reported *S. mansoni* glutamate receptors [21], where Smp_052660 contained a relatively well-conserved LBD with Smp_062660 appearing more atypical. Since all three *F. hepatica* glutamate GPCRs bear atypical LBDs with respect to both GABA_B_ and mGluR, it remains difficult to unequivocally define their ligand selectivity on the basis of conserved motifs. Nevertheless, the lack of *in silico* evidence for *F. hepatica* GABA_B_ GPCRs reflects the dominance of GABA_A_-like pharmacology, which suggests that flatworm GABA signal transduction is probably entirely mediated by ionotropic receptors [48,68].

**Fig 6:**
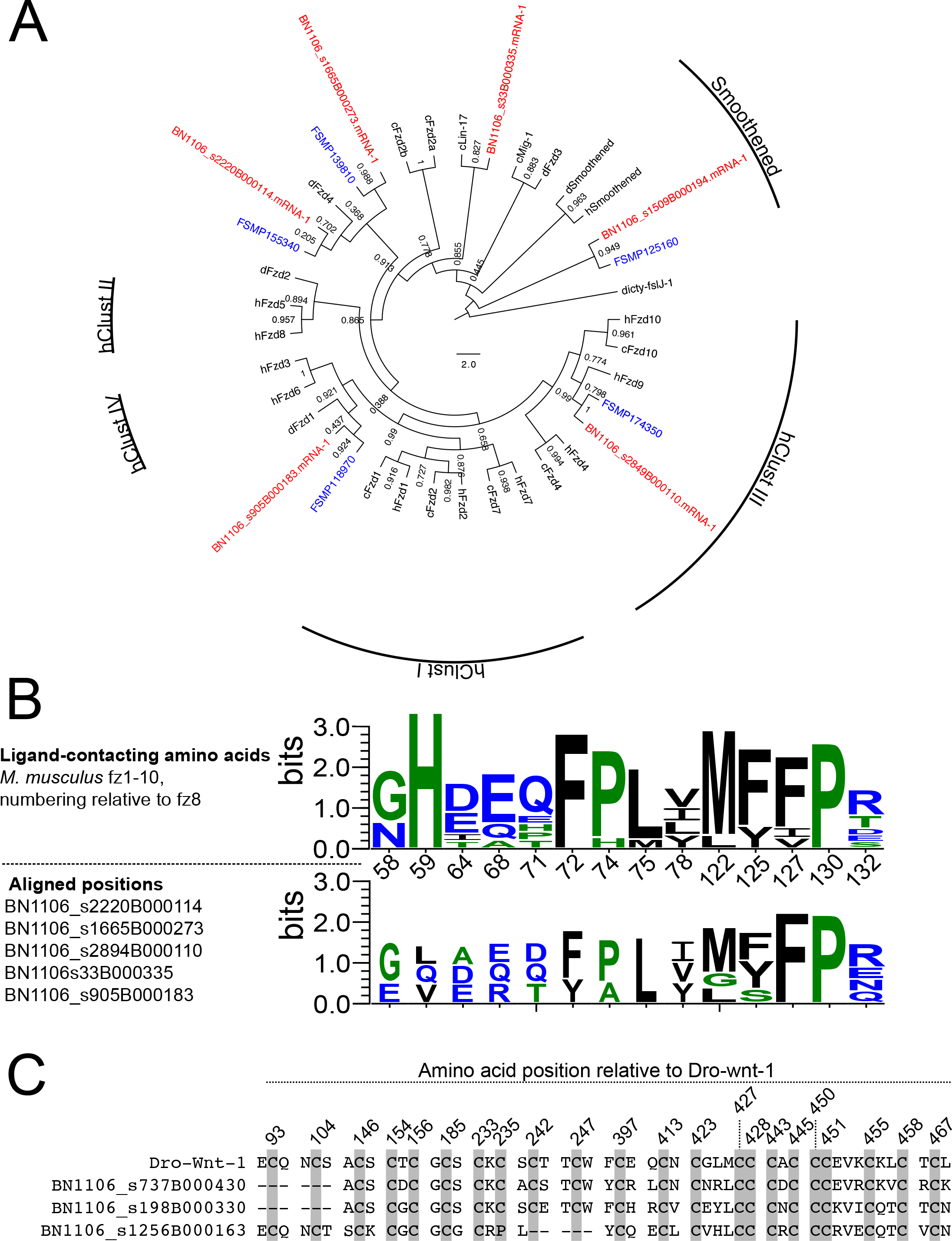
Frizzled/smoothened seven transmembrane receptors and wnt ligands in *Fasciola hepatica*. (A) Maximum likelihood phylogeny containing six *F. hepatica* frizzled/smoothened receptors, alongside those from *Schistosoma mansoni, Drosophila melanogaster, Caenorhabditis elegans* and *Homo sapiens* (identified by FSMP, d, c and h, respectively). *F. hepatica* sequences are coloured red, *S. mansoni* are coloured blue, all other species are coloured black. Radial labels indicate human frizzled clusters (hClust) I-IV, and the smoothened clade. Node numbers indicate statistical support as determined by approximate likelihood ratio test (aLRT). Tree was rooted against a *Dictyostelium* frizzled sequence (dicty-fslJ-1). Tree composition adapted from [21]. (B) WebLogo comparison of ligand interacting residues between mouse fz1-10 (top panel) and *F. hepatica* frizzled receptors. Numbering in top panel x-axis is relative to mouse fz8 [69]. (C) Three wnt-like sequences exist in *F. hepatica*. Shading indicates positions of 22 characteristic Cys residues, positions numbered relative to *D. melanogaster* wnt-1 (Dro-wnt-1).

### The Wnt binding domain is conserved in *F. hepatica* frizzled/smoothened receptors

Ten frizzled (fzd) GPCRs and a single smoothened (smo) GPCR are recognised in the human genome. In *F. hepatica* we identified five fzd-like sequences and one smo-like sequence (Figure 6; Table S1; Table S7). All of these show high scoring similarity to annotated sequences in the ncbi nr dataset (median E=3.8e^−83^), and all five fzd contain InterPro domain IPR000539, with the single smo containing domain IPR026544 (Table S1). Phylogenetic analysis of these alongside vertebrate and invertebrate receptors placed all in close proximity to existing fzd/smo groups (Figure 6A). Four *F. hepatica* fzd had individual direct orthologs with the four known *S. mansoni* fzd [21].

*Frizzled* receptors are activated by cysteine-rich glycoprotein ligands known as Wnts (Wingless and Int-1), and are involved in developmental signalling through at least three different signalling pathways [69]. Crystallography of mouse fz8, docked with *Xenopus* wnt8, identified 14 amino acids within the fz8 CRD that make contact with the Wnt8 ligand [69]. Positional conservation of these residues is apparent when fz8 is aligned with the five *F. hepatica* fzd sequences (Figure 6B; S6 Table), suggesting conservation of the wnt-frizzled interaction between liver fluke and vertebrates.

Two Wnt ligands have been described in *S. mansoni* [70,71]; our BLAST searches identified at least three Wnt-like sequences in the *F. hepatica* genome (BN1106_s198B000330.mRNA-1, BN1106_s1256B000163.mRNA-1, BN1106_s737B000430.mRNA-1; Figure 6C). These showed conservation of the 23 conserved cysteine residues that are diagnostic of Wnt glycoproteins [72]. Norrin, a non-Wnt protein ligand, can also activate Fz4, and the canonical β-catenin pathway. The amino acids involved in norrin binding to the fz4 CRD have also been determined [73], but we did not observe conservation of these in any of the *F. hepatica* fzd. Similarly, BLASTp searches of human norrin (Uniprot Q00604) against the *F. hepatica* genome did not return significant hits, suggesting that the norrin-fz signaling axis may not function in liver fluke. Smoothened receptors are structurally similar to frizzleds, but operate in a ligand-independent fashion within hedgehog signaling pathways that control several developmental processes [74]. Model organism genomes typically contain only one smoothened gene (SMO); this was the case in *S. mansoni* and *S. mediterranea* [21], and here we have identified a single *F. hepatica* smoothened (BN1106_s1509B000194; Figure 6; Table S1).

Fzd/smo GPCRs are involved broadly in the control of cellular development. Our discovery of fzd/smo GPCRs, and their Wnt ligands, in *F. hepatica* opens avenues towards probing molecular aspects of development and differentiation in the putative stem cells/neoblasts of liver fluke [35]. Neoblasts are the cells that impart the regenerative capacity of free-living turbellarian flatworms [75], and neoblast-like cells also represent the only proliferating cells in several parasitic species [76–78].

Therefore, these cells are important in understanding fundamental fluke biology and represent potential repositories of unique anthelmintic targets, capable of inhibiting worm growth or development. The presence of both receptor and ligand sequences will permit functional genomic dissection of Wnt-Frizzled ligand-receptor signalling networks, aimed at elucidating their roles in the development and differentiation of liver fluke neoblast-like cells. These FhGPCRs will enable comparisons between the biology of parasitic and free-living flatworms, where Wnt signaling is known to be essential for anterior-posterior polarity in regenerating planaria [79,80].

### Class B (Adhesion and Secretin) receptors

Class B receptors incorporate both adhesions and secretins. Adhesions are characterised by a long N-terminal extracellular domain (ECD) that includes several functional motifs. These ECDs are auto-proteolytically cleaved into two subunits that subsequently reassemble into a functional dimer [26]. We identified two Class B sequences in the *F. hepatica* genome (S7 Figure, S1 Table), both of which (scaffold181_78723-79604, and BN1106_s537B000355) contained GPCR class B InterPro domain IPR000832 and displayed closest BLASTp similarity (E=5.6e^−7^) to latrophilin-like receptors. These data suggest that both are adhesions, rather than secretins. Phylogenetic analysis of these GPCRs alongside human Class B receptors supports the definition of scaffold181_78723-79604 as an adhesion, alongside two previously reported *S. mansoni* adhesions (Smp_176830, Smp_099670) [21], while the other receptor appears more divergent. BN1106_s536B000355 pairs with another known *S. mansoni* adhesion (Smp_058380) [21], although both sit in closer proximity to human secretins than adhesions.

Deorphanization of a handful of adhesions matches them with a complex assortment of ligands including collagen, transmembrane glycoproteins, complement proteins and FMRFamide-like neuropeptides [81]. This assortment of potential ligands, and their expression in almost every organ system has led to the proposal of a diverse range of functions for vertebrate adhesions. The *F. hepatica* adhesion complement of two GPCRs is greatly reduced compared to the 33 receptors known in humans; in other flatworms 14, 4 and 1 adhesions have been described in *S. mediterranea, E. multilocularis* and *S. mansoni*, respectively [21,22,24]. Functional characterisation will be a challenging task given the wide range of possible functions to be assayed; an appealing starting point would be to investigate roles in neoblast motility prior to differentiation, given that mammalian adhesion GPCRs are involved in the control of cellular migration [81].

### Developmental expression

Using RNA-Seq methods, we were able to confirm expression of 101 GPCRs across libraries representing several *F. hepatica* life-stages. These datasets included publically available reads from individual developmental stages [18], and a transcriptome that we generated in-house for 21-day liver stage ex-vivo juveniles (juv2). Since these datasets were generated independently and clearly display distinct sequence diversities, we avoided any further direct comparisons between Cwiklinski juv1 and our juv2 datasets. Each dataset is analysed separately, below.

Figure 7A illustrates detection of 83 GPCRs across Cwiklinski’s developmentally staged RNA-Seq datasets. These comprised four FZD, thirteen aminergic rhodopsins, two opsins, 41 peptidergic rhodopsins, and 23 orphan rhodopsins. The latter included nine fwRhods. Clustering within Figure 7A’s expression heatmap shows clear developmental regulation of GPCR expression, outlining nine GPCRs with relatively higher expression in adults, two with higher expression in 21d juveniles, 64 GPCRs preferentially expressed in either 1h, 3h or 24h NEJs, and six receptors expressed most highly in eggs. GPCR classes appear to be randomly distributed across these expression clusters, giving little opportunity to infer function from expression. Adult-expressed GPCRs include five orphan fwRhods, three peptide receptors including a putative NPF/Y receptor, and a predicted octopamine-gated aminergic rhodopsin. The majority of expressed GPCRs occurred in the NEJ-focused expression cluster. Given data implicating GPCRs in motility, growth/development and sensory perception [11], it is no surprise to find high levels of GPCR expression in the NEJs, which must navigate and burrow their way from the gut lumen into the liver parenchyma, while also sustaining rapid growth from the start of the infection process. The high expression in these stages, of receptors that we predict to be activated by myomodulators such as ACh, FMRFamide, GYIRFamide, myomodulin, myosuppressin and 5-HT, provide tentative support for these predictions. The focused expression of six GPCRs in eggs suggests potential roles in the control of cellular proliferation and fate determination processes that occur during embryonation of liver fluke eggs. This complement did not include frizzled or adhesion GPCRs that are traditionally implicated in the control of development, instead consisting of rhodopsins (including an angiotensin-like peptide receptor, two octopamine-like amine receptors, one opsin receptor and one fwRhod receptor).

**Fig 7:**
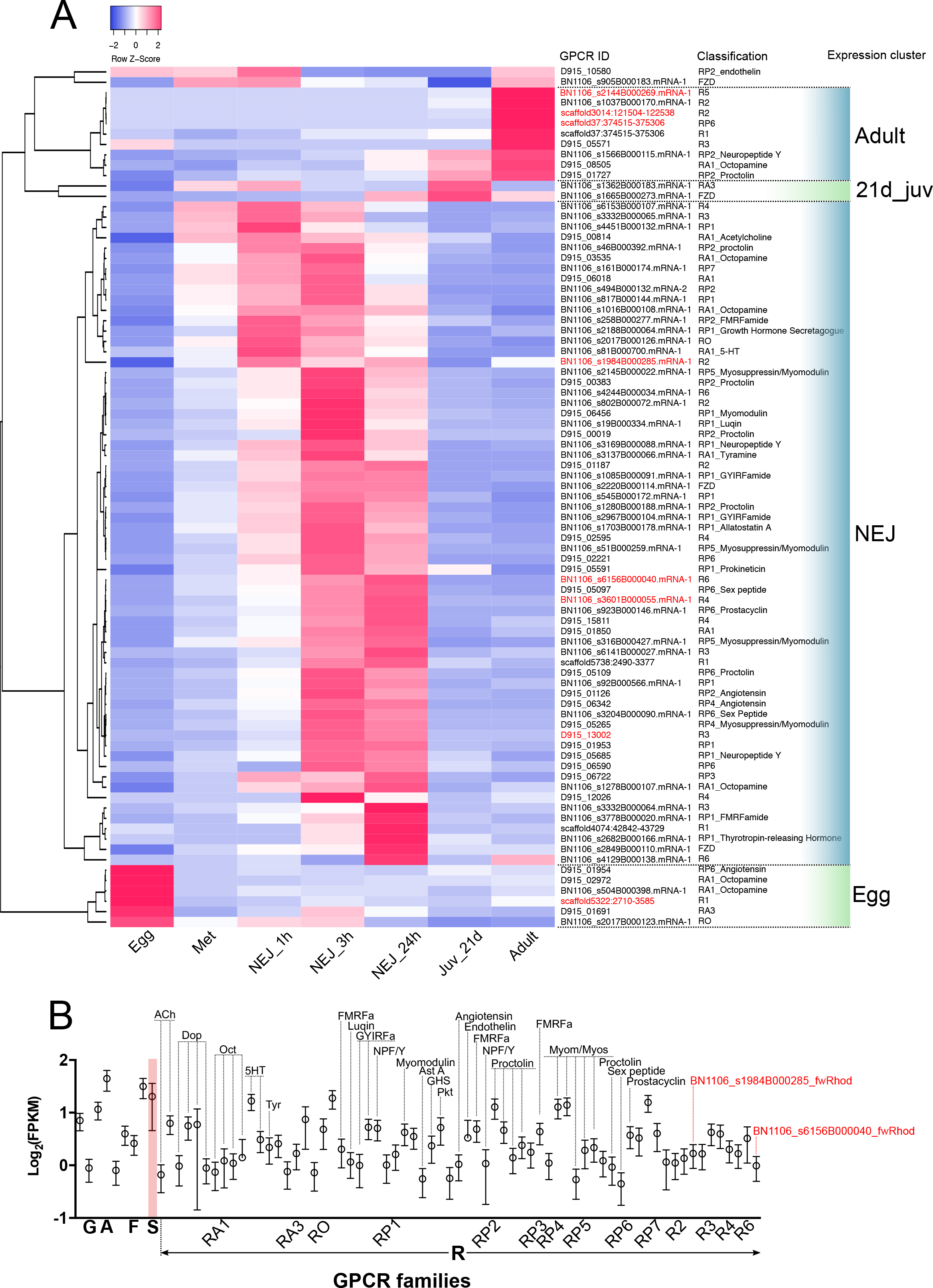
Expression profiling of 101 G protein-coupled receptors (GPCRs) in *Fasciola hepatica* life stages. (A) Expression heatmap generated from log_2_ FPKM values of 83 GPCRs identified from developmentally staged RNA-seq libraries. Life stages are represented in columns (Egg; Met, metacercariae; NEJ_1h, newly-excysted juvenile collected 1h post excystment; NEJ_3h, NEJ collected 3h post-excystment; NEJ_24h, NEJ collected 24h post-excystment; Juv_21d, liver stage juvenile parasites collected from murine livers 21 days following oral administration of metacercariae; Adult, adult parasites collected from the bile ducts of bovine livers). Rows indicate individual GPCRs, as denoted by the ID and phylogeny columns. The latter indicates receptor classification and predicted ligand where available (see S1 Table). Expression cluster column indicates clusters of GPCRs with highest expression focused in particular life stages. (B) Detection of 76 GPCRs in Illumina RNA-Seq libraries generated from *F. hepatica* 21 day liver-stage juveniles, recovered *ex vivo* from rat infections. Data show expression of three glutamate (G), one adhesion (A), four frizzled (F), one smoothened (S) and 67 rhodopsin (R) GPCRs. The rhodopsins include representatives of amine (RA1, RA3), opsin (RO), peptide (RP1-7), and orphan (R2,3,4,6). Data points (each at *n*=3) represent mean log_2_ FPKM ± 95% confidence intervals, as calculated by cuffdiff. In both panels, flatworm rhodopsins (fwRhods) are marked in red text. ACh, acetylcholine; AstA. Allatostatin A; Dop, dopamine; FMRFa, FMRFamide; GHS, growth hormone secretagogue; GYIRFa, GYIRFamide; Myom, myomodulin; Myos, myosuppressin; NPF/Y, neuropeptide F/Y; Oct, octopamine; Pkt, prokineticin; Tyr, tyramine; 5HT, 5-hydroxytryptamine.

Focusing on the pathogenic 21-day juvenile stage, we detected 76 GPCRs in our juv2 datasets, and 29 in the corresponding juv1 samples from Cwiklinski’s dataset (Figure 7B). Our juv2 dataset included three glutamate, one adhesion, four frizzled, one smoothened, and 67 rhodopsins. The identity of the receptors expressed here again attest to the key role of neuromuscular co-ordination in this highly motile life stage, which must penetrate and migrate through the liver parenchyma *en route* to the bile ducts. Amongst the receptors expressed in this stage and thought to have a role in neuromuscular function are several activated by classical neurotransmitters including ACh, dopamine and 5-HT. The peptide receptors include some with phylogenetic similarity to receptors for myoactive flatworm peptides (FMRFamide, GYIRFamide, NPF) [11], as well as receptors from other invertebrates activated by peptide ligands known to have excitatory effects on flatworms (allatostatin A, myomodulin, proctolin) [82]. The presence of highly expressed GPCRs with probable neuromuscular functions in liver stage juveniles, points to the importance of studying these receptors with a view to flukicide discovery. The damage caused by migrating juvenile fluke requires that new flukicides are effective against this stage. The neuromuscular GPCRs expressed in migrating juveniles provide compelling targets for new drugs.

### Conclusions

GPCRs are targets for 33% of human pharmaceuticals [17], illustrating the appeal of GPCRs as putative anthelmintic targets. This study provides the first description of the *F. hepatica* GPCR complement permitting consideration of a GPCR target-based screening approach to flukicide discovery. To facilitate the deorphanization experiments that will precede compound screening efforts, we have described a set of high confidence rhodopsin ligand-receptor pairs. We identified these GPCRs, including receptors for ACh, octopamine, 5HT and NPF/Y, through phylogenetic comparison with existing deorphanised receptors and positional conservation of ligand-interacting residues within ligand binding domains. Our additional descriptions of flatworm-specific rhodopsins support the potential for synthetic ligands to be parasite-selective anthelmintics.

## Materials and Methods

### Liver fluke sequence databases

We exploited two *F. hepatica* genome assemblies available from WormBase ParaSite [83], generated by Liverpool University (http://parasite.wormbase.org/Fasciola_hepatica_prjeb6687/Info/Index/; [18], and Washington University, St Louis (http://parasite.wormbase.org/Fasciola_hepatica_prjna179522/Info/Index/; [19].

### Identification of GPCR-like sequences from *F. hepatica*

Figure 1 summarises our GPCR discovery methodology, which employed Hidden Markov Models (HMMs) constructed from protein multiple sequence alignments (MSAs) of previously described *S. mansoni* and *S. mediterranea* GPCR sequences [21]. Individual HMMs were constructed for each GRAFS family [25]. Alignments were generated in Mega v7 (http://www.megasoftware.net) [84] using the Muscle algorithm with default parameters. HMMER v3 (http://.hmmer.org) was employed to construct family-specific HMMs (*hmmbuild*) from alignments and these were searched (*hmmsearch*) against a predicted protein dataset from *F. hepatica* genome PRJEB6687 consisting of 33,454 sequences [18]. Returned sequences were filtered for duplicates and ordered relative to the *hmmsearch* scoring system, enabling the classification of hits according to the GRAFS family to which they showed most similarity (i.e. highest score, lowest E value). All remaining returns were then used as BLAST queries (BLASTp and tBLASTn) to identify matching, or additional, sequences originating from the PRJEB6687 and PRJNA179522 genomes (Figure 1). Where sequences appeared in both genomes, we kept the longest annotated sequence (S1 Table).

### GPCR annotation

Sequences resulting from HMM searches were filtered by transmembrane (TM) domain composition, using hmmtop (http://www.sacs.ucsf.edu/cgi-bin/hmmtop.py) [85,86]. Sequences containing ≥4 TMs were analysed as described below.

### Homology analyses

All GPCRs were used as BLASTp [87] queries, to identify their closest (highest scoring) match in the ncbi non-redundant (nr) protein sequence dataset (https://blast.ncbi.nlm.nih.gov/Blast.cgi), with default settings and the “Organism” field set to exclude Platyhelminthes (taxid: 6157). All GPCRs were additionally searched against more phylogenetically limited datasets, by using the “Organism” field to limit the BLASTp searches to: (i) Basal phyla, Ctenophora (taxid:10197), Porifera (taxid:6040), Placozoa (taxid:10226), Cnidaria (taxid:6073); (ii) Superphylum Lophotrochozoa (taxid: 1206795), excluding phylum Platyhelminthes (taxid: 6157); (iii) Superphylum Ecdysozoa (taxid: 1206794); (iv) Superphylum Deuterostomia (taxid: 33511). For BLASTp searches against other flatworms, we performed local BLAST+ [88] on the WBPS9 release of WormBase Parasite, which included predicted protein datasets from 30 flatworm species. In all cases, we recorded the single highest scoring hit, or recorded “no significant similarity found” in cases where no hits were returned (Table S1); sequences generating both GPCR hits and “no hits” were retained. Where the top hit was not to a GPCR, that sequence was removed from the dataset.

### Domain composition

GPCR identities were confirmed using InterProScan Sequence Search (http://www.ebi.ac.uk/interpro/search/sequence-search) [89] and/or HMMER HMMScan (http://www.ebi.ac.uk/Tools/hmmer/search/hmmscan) [90]. Again, sequences returning non-GPCR domains were omitted from the dataset, with all others retained.

### Motif identification

As an additional measure of confidence in our identifications, we analysed the presence/absence of key motifs diagnostic of receptor families and subfamilies. These analyses were performed for rhodopsins generally, the ligand binding domains (LBDs) of rhodopsin receptors for acetylcholine (ACh), neuropeptide F/Y (NPF/Y), octopamine and serotonin (5-hydroxytryptamine, 5HT), and for the LBDs of glutamate and frizzled/smoothened families. Motifs were identified via protein multiple sequence alignment (MSA) of GPCRs, performed in MAFFT (http://www.mafft.cbrc.jp/alignment/server) [91]. Only identical amino acids were accepted at each site, with conservation expressed as % identity across all sites. Motif illustrations (Figures 3 & 7) were generated using WebLogo 3 (http://weblogo.threeplusone.com) [92].

### Phylogenetic reconstruction

Maximum likelihood (ML) phylogenetic trees were constructed using PhyML (http://www.phylogeny.fr) [93], from protein MSA generated in MAFFT (http://www.mafft.cbrc.jp/alignment/server/) [91]. Alignments were manually edited (in Mega v7) to include only TM domains, by removing extramembrane blocks aligned with human glutamate, rhodopsin, adhesion or frizzled proteins. Trees were constructed from these TM-focused alignments in PhyML using default parameters, with branch support assessment using the approximate likelihood ratio test (aLRT), under “SH-like” parameters. Trees, exported from PhyML in newick format were drawn and annotated in FigTree v1.4.2 (http://tree.bio.ed.ac.uk/software/figtree/).

### RNA-Seq analyses

Expression of *F. hepatica* GPCRs was investigated in publically available and in-house generated RNA-Seq datasets. These included developmentally staged Illumina transcriptome reads associated with one of the *F. hepatica* genome projects [18] (reads accessed from the European Nucleotide Archive at http://www.ebi.ac.uk/ena/data/search?query=PRJEB6904). These samples originated from distinct developmental stages of US Pacific Northwest Wild Strain *F. hepatica* (Baldwin Aquatics), including egg (*n*=2), metacercariae (met; *n*=4), *in vitro* NEJs 1h post-excystment (NEJ1h; *n*=1), *in vitro* NEJs 3h post-excystment (NEJ3h; *n*=2), *in vitro* NEJs 24h post-excystment (NEJ24h; *n*=2), *ex-vivo* liver-stage juveniles (juv1; *n*=1) and *ex vivo* adult parasites (Ad; n=3). Our in-house datasets were generated from *ex vivo* liver stage *F. hepatica* juveniles (Italian strain, Ridgeway Research Ltd, UK), recovered from rat (Sprague Dawley) hosts at 21 days following oral administration of metacercariae (juv2; *n*=3). Total RNA, extracted with Trizol (ThermoFisher Scientific) from each of the 3 independent biological replicates, was quantified and quality checked on an Agilent Bioanalyzer, converted into paired-end sequencing libraries and sequenced on an Illumina HiSeq2000 by the Centre for Genomic Research at the University of Liverpool, UK. RNA samples were spiked prior to library construction with the ERCC RNA Spike-In Mix (ThermoFisher Scientific) [94]. All read samples were analysed using the TopHat and Cufflinks pipeline [95–100], with mapping against PRJEB6687 genome sequence and annotation files (accessed from WormBase Parasite; http://parasite.wormbase.org/ftp.html). Data were expressed as number of fragments mapped per million mapped reads per kilobase of exon model (FPKM). In juv2 datasets we discarded GPCRs represented by fewer than 0.5 FPKM (the minimum linear sensitivity that we detected with our ERCC spike in); for the staged datasets, we included only receptors represented by ≥0.5 FPKM in at least one life stage. Heatmaps were generated with heatmapper (http://www.heatmapper.ca/) [101] set for Average Linkage, and Pearson Distance Measurement.

---

**S1 Table. *Fasciola hepatica* G protein coupled receptor (GPCR) dataset summary**. Table describes sequences containing 4-9 transmembrane (TM) domains. Each sequence is defined in terms of GRAFS family, and annotated for TM composition, sequence length, phylogeny, domain composition and homology relative to various datasets.

**S2 Text. *Fasciola hepatica* G protein-coupled receptor (GPCR) protein sequence dataset**.

**S3 Table. Flatworm-specific rhodopsins (fwRhods) in *Fasciola hepatica* and other flatworms**. Species and genome IDs of sequences that have ≥4 transmembrane (TM) domains, and lack high-scoring orthologues in non-flatworms (BLASTp score E≥0.01 vs ncbi nr excluding Platyhelminthes), and show conservation of at least two of the four ubiquitous rhodopsin motifs. Each sequence is annotated for protein domains where present (Pfam HMMScan). Accessions refer to WormBase ParaSite.

**S4 Table. Rhodopsin ubiquitous motifs and ligand binding domains for ACh, NPF/Y 5-HT, octopamine**. Note data are in individual tabs. Rhodopsin: Sequence motifs extracted from alignment of *F. hepatica* rhodopsins, corresponding to ubiquitous rhodopsin motifs of TMs 2, 3, 6 and 7; Acetylcholine, NPF, 5-HT, Octopamine: Amino acids extracted from alignment of *F. hepatica* rhodopsins with mutationally or structurally-characterised GPCRs (comparators). Summary: Percent identity of ACh, NPF, 5-HT and Oct receptor LBDs, indicating most conserved LBD sequences showing at least 75% identity.

**S5 Fig. Phylogenetic comparison of *Fasciola hepatica* GPCRs with deorphanised bilaterian GPCRs**. (A) Peptide receptors (*F. hepatica* black or magenta as described in Fig x); (B) Amine receptors (*F. hepatica* dark blue); In all cases, non-flatworm receptors are coloured light blue. In (A) outer labels indicate positions of receptors for neuropeptide families previously reported in flatworms (McVeigh et al., 2009; Collins et al., 2010; Koziol et al., 2016); in (B), outer labels represent major groups containing phylogenetically similar *F. hepatica* sequences. Trees were midpoint rooted, maximum likelihood phylogenies of transmembrane domains I-VII. Numbers at nodes indicate statistical support from approximate likelihood ratio test (aLRT). Scale bars at the centre of each tree indicate number of substitutions per site. Abbreviations: ACh, acetylcholine; CCAP, crustacean cardioactive peptide; Dop, dopamine; FLP, FMRFamide-like peptide; GrH, gonadotropin-releasing hormone; Luq, luqin; Mmd, myomodulin; NKY, neuropeptide KY; NPF/Y, neuropeptide F/Y; Oct, octopamine; PK, pyrokinin; SIFa, SIFamide; Tyr, tyramine; SmGPR, schistosome GPCRs; 5HT, 5-hydroxytryptamine.

**S6 Table. Frizzled receptor ligand binding domain motifs**. Amino acids extracted from alignment of *F. hepatica* frizzled and smoothened GPCRs with mutationally characterised mouse fz8, also showing positionally-conserved residues in mouse fz1-10 (top panel) and *F. hepatica* frizzled receptors. Numbering in top row relative to mouse fz8 (Janda et al., 2012). Green boxes indicate identical residues in *F. hepatica* vs mammalian Fzd.

**S7 Figure: Adhesion receptor phylogeny**. Maximum likelihood phylogeny of *Fasciola hepatica* adhesion/secretin-like GPCRs alongside class B GPCRs from human and *Schistosoma mansoni*. Tree was a midpoint rooted, maximum likelihood phylogeny of transmembrane domains I-VII. Numbers at nodes indicate statistical support from approximate likelihood ratio test (aLRT). Scale bars at the centre of each tree indicate number of substitutions per site.

